# Vinculin Y822 phosphorylation regulates cardiomyocyte adhesion dynamics and adherens junction maturation in the heart

**DOI:** 10.1101/2024.10.28.620745

**Authors:** Xiaofei Li, Rainy Wortelboer, Yi Song, Sahana Balasubramanian, Callie McLain, Alex Hernandez Manriquez, Joseph Suh, Brenton D. Hoffman, Adam V. Kwiatkowski, Glenn L. Radice

## Abstract

In the heart, cell-matrix and cell-cell adhesions reorganize in response to increased cardiac demand and growth to promote cardiomyocyte maturation. Vinculin, a mechanosensitive adaptor protein, links filamentous actin to cell-matrix and cell-cell adhesions and is thus positioned to regulate adhesion reorganization. However, how the two adhesion systems are coordinated in the heart, and the role of vinculin in this process is poorly understood. Here, we define the role of vinculin phosphorylation at tyrosine residue 822 (pY822) in cardiomyocyte adhesion and heart function. We found that pY822 correlated with dynamic junction remodeling in the developing heart but was lost as junctions matured postnatally. We then mutated Y822 to phenylalanine (Y822F) in the mouse to determine pY822 function *in vivo*. Homozygous mutant *Vcl Y822F* mice were viable and exhibited normal cardiac function at ten weeks of age; however, cardiac dysfunction was observed at 28 weeks. Vinculin and adherens junction proteins were reduced at cardiomyocyte junctions in *Y822F* hearts. In contrast, α5/β1 integrin and fibronectin increased along the lateral border of *Y822F* cardiomyocytes. Our results demonstrate that vinculin Y822 phosphorylation regulates the balance between cadherin and integrin adhesion organization, highlighting the importance of post-translational modification in modulating vinculin function in heart physiology.

## Introduction

Mechanical forces are critical in regulating cellular behavior and function in all cell types. The ability of a cell to sense, integrate, and convert mechanical stimuli into biochemical signals to mediate intracellular changes is called mechanotransduction. Defects in mechanotransduction are implicated in the pathogenesis of various diseases, including arteriosclerosis, cardiomyopathies, asthma, and cancer (1–4). The ability to detect and respond to changes in mechanical stimuli is essential in the heart as it allows the cardiomyocyte (CM) to react and adapt to increased mechanical load during cardiac development and disease. For optimal force transmission, CMs must maintain physical linkages with 1) neighboring cells, mediated by cadherin-based cell-cell adhesion complexes, and 2) the surrounding extracellular matrix (ECM), mediated by integrin-based cell-ECM adhesion complexes. Multiple cytoskeletal adaptor proteins function together to transduce mechanical stimuli across cell-cell and cell-ECM adhesion complexes. Among these, vinculin (VCL) is critical in linking integrin- and cadherin-based adhesions to the actin cytoskeleton. It is thus well-positioned to coordinate mechanical responses in CMs.

VCL regulates adhesion dynamics by directly binding to filamentous actin (F-actin), stimulating actin polymerization, and recruiting actin-remodeling proteins (5–7). Structurally, VCL comprises a series of helical bundles organized into five domains, termed D1-D5. D1-D4 are a collection of seven four-helix bundles (the VCL “head-domain”) that mediate binding to α-catenin, talin, and other ligands. D5 is a five-helix bundle (the VCL “tail-domain”) that binds F-actin. VCL exists primarily in two conformations in the cell: an “open,” active form and a closed, auto-inhibited state in which the N-terminal head domain (D1-D4) forms extensive interactions with the C-terminal tail (D5) (6). Relieving VCL auto-inhibition to permit F-actin binding requires binding to α-catenin at adherens junctions (AJs) and talin at focal adhesions (FAs), known as costameres in muscle cells (8, 9). VCL is subject to extensive biochemical regulation. Phosphorylation of multiple residues, e.g., Y100, Y1065, S1033, S1054, and Y822, regulates VCL activity at FAs and AJs (6, 10). Y822 is in VCL D4 adjacent to the linker region that connects the head and tail domains and is phosphorylated in response to mechanical force by the non-receptor tyrosine kinase Abl1 (11, 12). Y822 phosphorylation promotes an “open”, active conformation that facilitates VCL binding to the cadherin/catenin complex and VCL-mediated recruitment of the actin nucleator VASP (11, 12). VASP binding could, in turn, induce actin assembly and increase tension at AJs (12). Biophysical studies using VCL FRET-based tension sensors have provided detailed molecular insight into VCL force transmission in cells grown in 2-D culture (8, 9, 13–17). Notably, mutations in Y822 have been linked to disease in humans. A *Vcl* mutation resulting in a cysteine substitution at Y822 was identified in uterine cancer (Cancer Genome Atlas Database) (18) and a patient with dilated cardiomyopathy (ClinVar, VCV002171895.1). In addition to blocking phosphorylation, Y822C VCL exhibits aberrant disulfide bond formation with the FA protein paxillin, and Y822C mutant cells have numerous small FAs that grow better in soft agar than control cells (18). The existence of disease-associated mutations in Y822 reinforces the importance of this residue in VCL biology.

Shortly after birth, CMs elongate, and myofibrils align during maturation to eventually form rod-shaped, terminally differentiated CMs connected end-to-end by a specialized junction called the intercalated disc (ICD) (19, 20). The ICD is formed from three junction types: AJs and desmosomes that connect the actin and intermediate filament cytoskeletons, respectively, of adjoining CMs, and gap junctions that electrochemically couple CMs. During CM maturation, N-cadherin, which is initially distributed around the entire cell border, becomes restricted to the bipolar ends of the cell. This polarized redistribution is required for ICD formation and efficient cardiac organization/function. Dynamic remodeling of CM-ECM interactions is also required for cardiac morphogenesis. For example, cardiac outflow tract formation depends on α5/β1 integrin-fibronectin (FN) adhesions (21). After birth, FN declines, and collagen and laminin increase along the lateral membrane, resulting in stiffer cardiac tissue. Corresponding shifts in integrin expression accompany changes in ECM. Notably, after a heart attack, CM cell-cell and cell-ECM interactions remodel in the border zone of infarcted myocardium (22) to resemble those seen in the neonatal heart (19). Whether VCL coordinates the interplay between the two cell adhesion structures in the heart remains unclear.

Evidence from human and animal studies has revealed a critical role for VCL in embryonic development, tissue homeostasis, and disease (23–26). VCL is essential for maintaining the structural integrity of heart muscle as mutations in human *VCL* have been associated with both dilated and hypertrophic forms of cardiomyopathy (27–29). In mice, global *Vcl* knockout results in mid-gestation lethality with multiple development abnormalities, including cardiac defects (24). Cardiac-specific loss of *Vcl* causes ICD defects, and half of the *Vcl* mutant mice exhibit sudden death (< 3 months), while the surviving mice develop dilated cardiomyopathy and die at six months of age (26). N-cadherin and β1 integrin are reduced in VCL-deficient adult CMs, explaining the severity of the mechanoelectrical coupling defects in the mutant mice. In cultured CMs, VCL is required to link myofibrils to nascent cell-cell contacts (30). Finally, VCL activation regulates myofibril maturation in the developing zebrafish heart in response to the mechanical forces generated by increasing cardiac contractility (31). However, if and how posttranslational modification regulates VCL function in cardiac physiology and disease remains unexplored.

Here, we show that pY822 VCL expression correlates with dynamic adhesion structures during cardiac morphogenesis and declines when AJs mature in the postnatal heart. To define the requirement for pY822 VCL *in vivo*, we created a new strain of *Vcl* mutant mice in which Y822 was mutated to a non-phosphorylatable phenylalanine (F) using CRISPR/Cas9-mediated gene editing. Loss of pY822 disrupted CM cell adhesion organization in the postnatal heart, notably reduced AJ maturation and enhanced integrin-fibronectin interactions. Defects in adhesion organization were also observed in cultured *Vcl Y822F* neonatal cardiomyocytes, corroborating the *in vivo* studies. *Vcl Y822F* mice exhibited cardiac dysfunction with age. These data provide the first genetic evidence that VCL phosphorylation at Y822 regulates CM adhesion dynamics and cardiac function.

## Results

### Decline in VCL Y822 phosphorylation is associated with AJ maturation in the postnatal cardiomyocyte

Phosphorylation sites have been mapped to all domains of VCL (6), but the site most frequently identified by mass spectrometry is amino acid Y822 near the end of D4. While *in vitro* and cell-based studies have provided insight into how pY822 regulates VCL activity, the functional significance of VCL pY822 on embryonic morphogenesis and tissue mechanics remains unexplored. We focused on the heart, which is subject to dynamic mechanical stresses throughout life. We first examined the expression profile of VCL pY822 in wild-type (C57BL/6J) heart tissue using a VCL pY822-specific antibody. Western blot analysis of heart lysates revealed that VCL pY822 was abundant in the embryonic/perinatal period but declined significantly by postnatal day (P) 14 and remained low in 10-week-old adult mice **(Fig. 1A, B)**. We then examined VCL pY822 localization by immunostaining. Heart sections from E13.5, P1, P14, and 10-week-old adult mice were stained for VCL pY822 and VCL **(Fig. 1C)**. VCL pY822 was predominantly nonjunctional, displaying a punctate and diffuse staining pattern in CMs. Note that two different monoclonal antibodies (mAbs) against VCL did not recognize the VCL pY822 population. Similarly, a mAb against VCL was reported not to recognize VCL pY1065 (32), suggesting the altered conformation of phosphorylated VCL is not recognized by the VCL mAbs. Thus, VCL pY822 expression correlates with dynamic adhesion structures and cell rearrangements during heart development and is lost upon the establishment of mature adhesive structures in the adult working myocardium.

**Figure 1.**
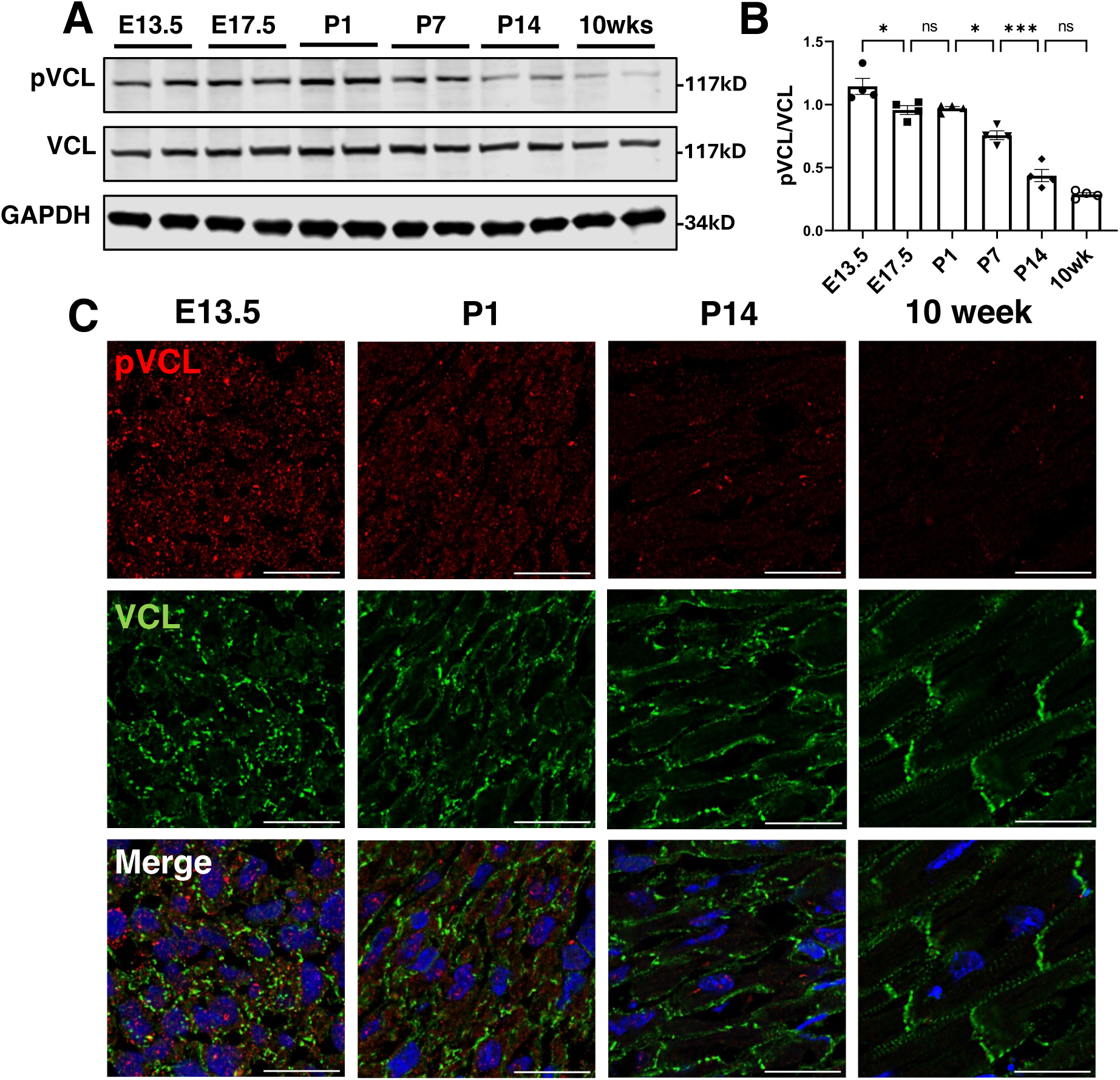
pY822 VCL expression during heart development. (A) Western blot analysis of heart lysates from embryonic (E), postnatal (P), and adult 10-week-old WT (C57BL/6J) mice. (B) Quantification of pVCL/VCL (n=4 hearts/time point). *, *p*<0.05, ***, *p*<0.001, ns, not significant by Students t-test (B) Quantification of western blot Y822 phosphorylated vinculin/vinculin intensity. *P<0.05, ***P<0.001 by two-tailed unpaired Student’s t-test. Error bars represent standard error of the mean (SEM). (C) Representative deconvolved max projection images of 2 slices with 0.175μm step size of heart sections from WT mice co-stained with pY822 VCL (red), VCL (green), and DAPI (blue). Scale bar: 20μm.

### VCL Y822 phosphorylation is not required for embryonic morphogenesis

To investigate the function of VCL pY822 *in vivo*, we created a new strain of *Vcl* knock-in mice using CRISPR/Cas9-mediated gene editing. *Vcl Y822* was mutated to a non-phosphorylatable phenylalanine (F) to generate *Vcl^Y822F/+^* mice **(Fig. 2A-C)**. In contrast to *Vcl-null* mice (24), *Vcl^Y822F/Y822F^* animals were born at the expected Mendelian frequency (53, *Vcl^+/+^*; 136, *Vcl^Y822F/+^*; 65, *Vcl^Y822F/Y822F^*, n=254) and showed no obvious macroscopic phenotypic abnormality. Loss of pY822 VCL in the *Vcl* mutant hearts was confirmed by immunoblotting and immunostaining **(Fig. 2D, F)**. Importantly, VCL Y822F protein levels in mutant mice were similar to WT VCL levels in control animals, as determined by immunoblot analysis of heart lysates **(Fig. 2D, E)**. Next, we examined pY822 expression in P1 heart sections. The punctate pY822 staining pattern was lost in *Vcl* mutant CMs **(Fig. 2F)**. We conclude that phosphorylation at VCL Y822 is not required for embryonic morphogenesis and viability.

**Figure 2.**
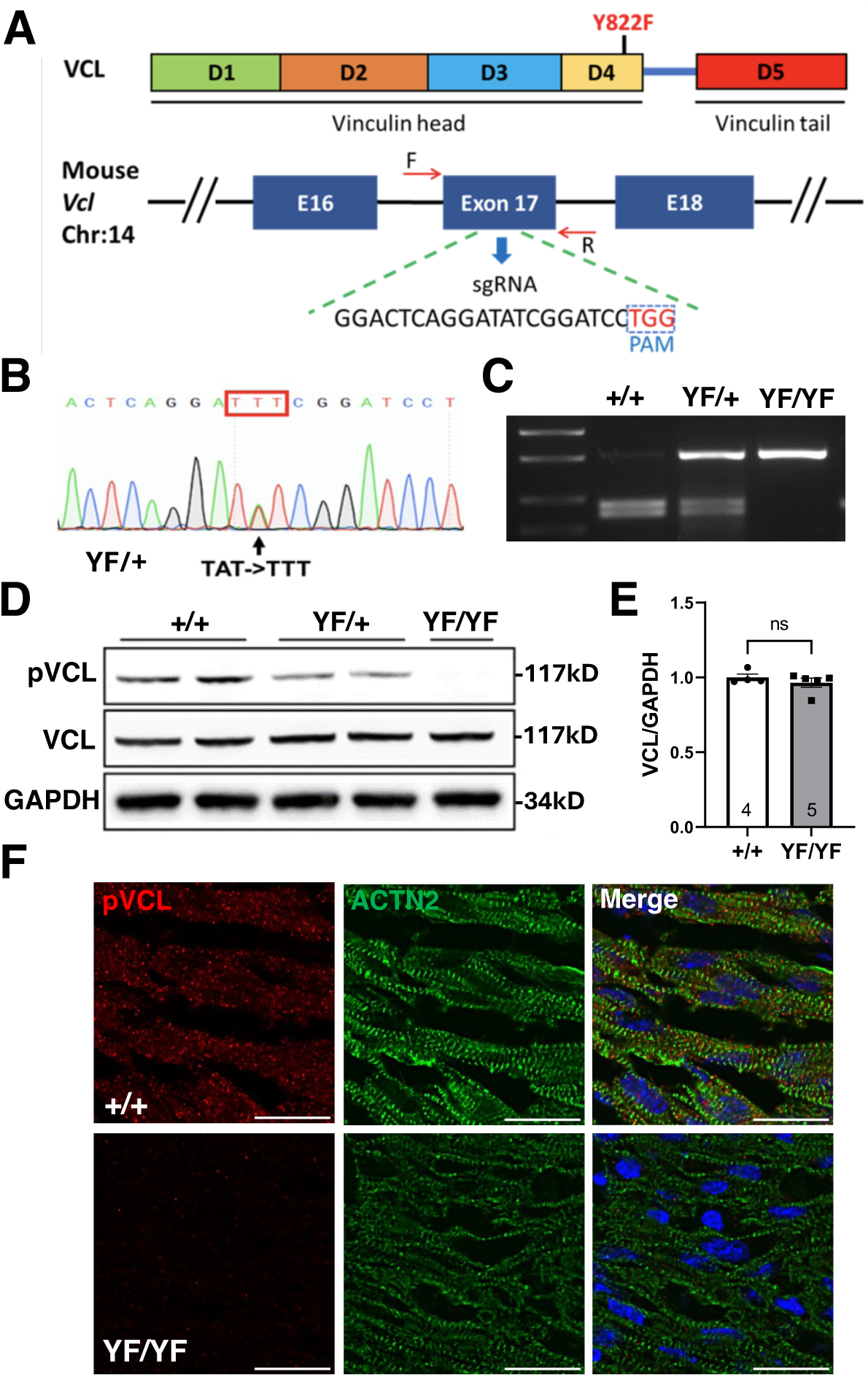
Generation of *Vcl Y822F* mutant mice. (A) Schematic of VCL protein structure indicating location of amino acid change and gene locus showing targeted exon with sgRNA target sequence. (B) Sanger sequencing shows the nucleotide change causing the *Vcl Y822F/+* (YF) mutation. (C) Genotyping of the mutant mice by PCR. The mutation destroyed an EcoRV site in exon 17 of the *Vcl* gene. The PCR product (514 bp) was digested with EcoRV resulting in two WT bands (275 and 239 bps) or undigested full-length band (514 bp) corresponding to the YF mutation. (D) Western blot analysis of WT, YF/+, and YF/YF P4 heart lysates. (E) Quantification of WT and VCL YF protein levels. (F) Representative immunofluorescence images of heart sections from WT and *Vcl Y822F* P1 mice co-stained for pY822 VCL (red) and α-actinin (green). Scale bar: 25μm.

### Loss of VCL pY822 alters the balance between cell-cell and cell-ECM adhesions in the postnatal heart

Shortly after birth, myocardial cell-cell and cell-ECM adhesions reorganize as cardiomyocytes mature in response to increased cardiac demand and growth. Thus, the postnatal heart offers a unique system to investigate junction remodeling and maturation. To probe effects on AJ development and organization, we examined junction maturation in P28 heart sections by staining for VCL and β-catenin **(Fig. 3A)**. Mutant VCL Y822F was reduced at CM termini (i.e., ICD) compared to the lateral membrane **(Fig. 3B)**. Also, β-catenin intensity was weaker at the ICD **(Fig. 3D)**, consistent with a reduction in AJs. Moreover, VCL and β-catenin appeared punctate in the *Vcl Y822F* mutant compared to the linear pattern observed in the WT heart **(Fig. 3A, inset)**. Next, we examined the expression of the N-cadherin/catenin complex in heart lysates from P14 and P28 mice **(Fig. S1)**. There was no change in protein levels of N-cadherin, αE-catenin, αT-catenin, and β-catenin between WT and *Vcl Y822F* hearts. Moreover, there was no change in the expression of afadin, a cytoskeletal adaptor protein associated with the cadherin/catenin complex **(Fig. S1)**.

**Figure 3.**
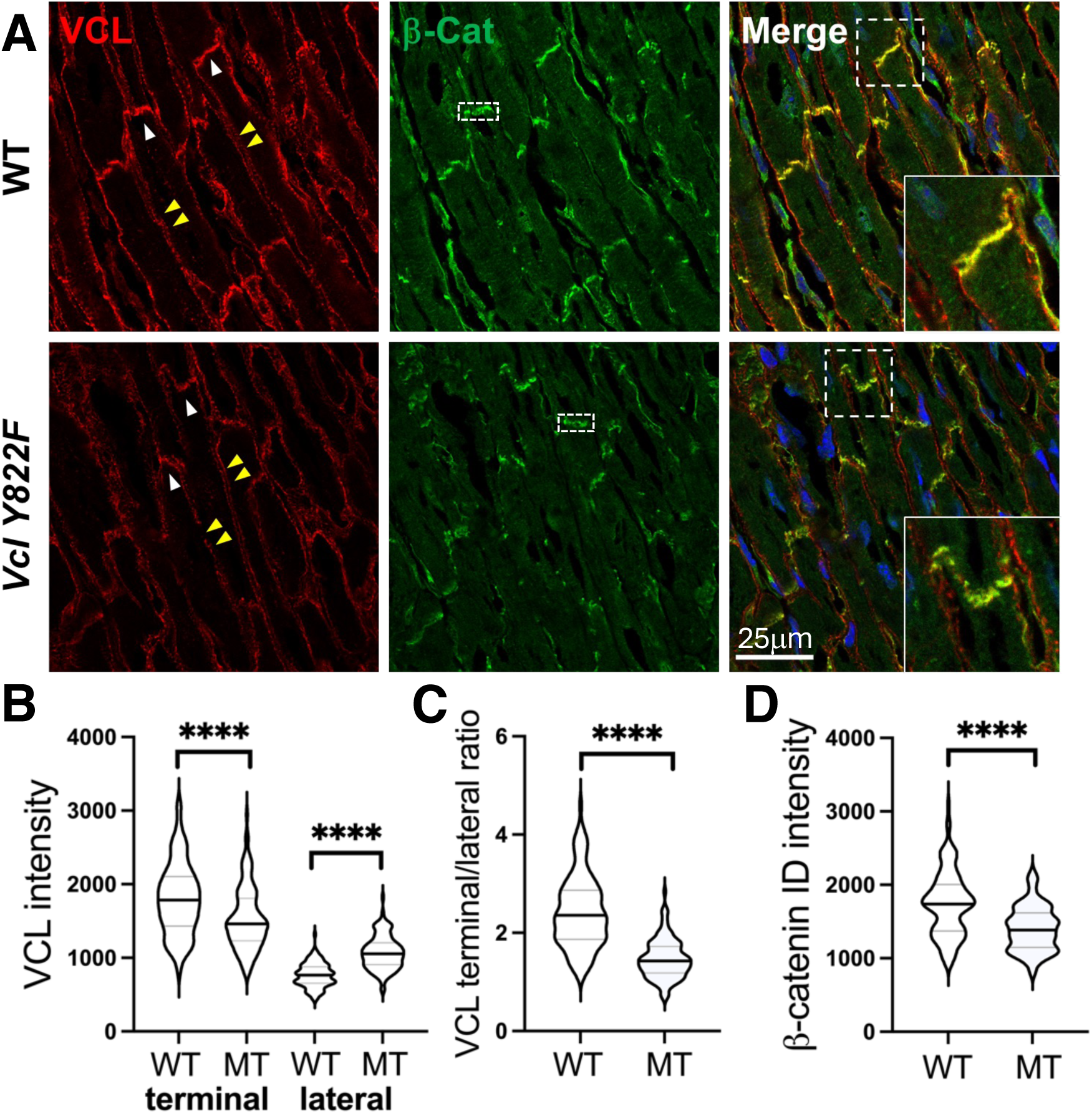
Blocking VCL phosphorylation leads to reduced N-cadherin adhesion complexes at the ICD. (A) Representative immunofluorescence images of P28 heart sections from WT and *Vcl* mutant (MT) co-stained with VCL (red) and β-catenin (green). Scale bar: 25μm. (B) Violin plots of CM VCL fluorescence intensity at terminal (white arrowhead) and lateral membrane (double yellow arrowheads). (C) Ratio of VCL terminal/lateral intensity. (D) Violin plots of β-catenin intensity at termini (stippled box). n=115 CMs from 4 hearts/genotype, ****, *P*<0.0001 by two-tailed unpaired Student’s t-test. Error bars represent SEM.

To probe the effects on FA and ECM organization, we examined the expression of fibronectin (FN) and its primary receptor α_5_β_1_ integrin in *Vcl Y822F* mutant and control hearts. Immunostaining of heart sections from P7 mice revealed a marked increase in α5 integrin and FN expression in *Vcl Y822F* mutants compared to controls **(Fig. 4A)**. Integrin-FN binding leads to the formation of FN fibrils that are initially soluble in the detergent deoxycholate (DOC) but are gradually converted into a stable, DOC-insoluble form that comprises the mature matrix (33). Both α5 and FN increased in the DOC-insoluble fraction, consistent with the strengthening of myocardial cell-matrix interactions in the *Vcl Y822F* hearts **(Fig. 4B)**. We then questioned if other VCL regulatory pathways were altered in *Vcl Y822F* mice. For example, phosphorylation of VCL Y100 and Y0165 by Src kinase regulates VCL protein dynamics at FAs (32, 34). Therefore, we examined VCL pY100 and pY1065 in *Vcl Y822F* hearts using phospho-specific antibodies. We observed no difference in pY100 or pY1065 levels in *Vcl Y822F* P7 heart lysates versus control **(Fig. S2)**, suggesting that Src-mediated signaling through VCL at FAs is not perturbed in the *Vcl Y822F* mutant. These results show that VCL Y822 phosphorylation coordinates mechanical homeostasis in CMs by regulating the balance between cadherin- and integrin-based adhesions.

**Figure 4.**
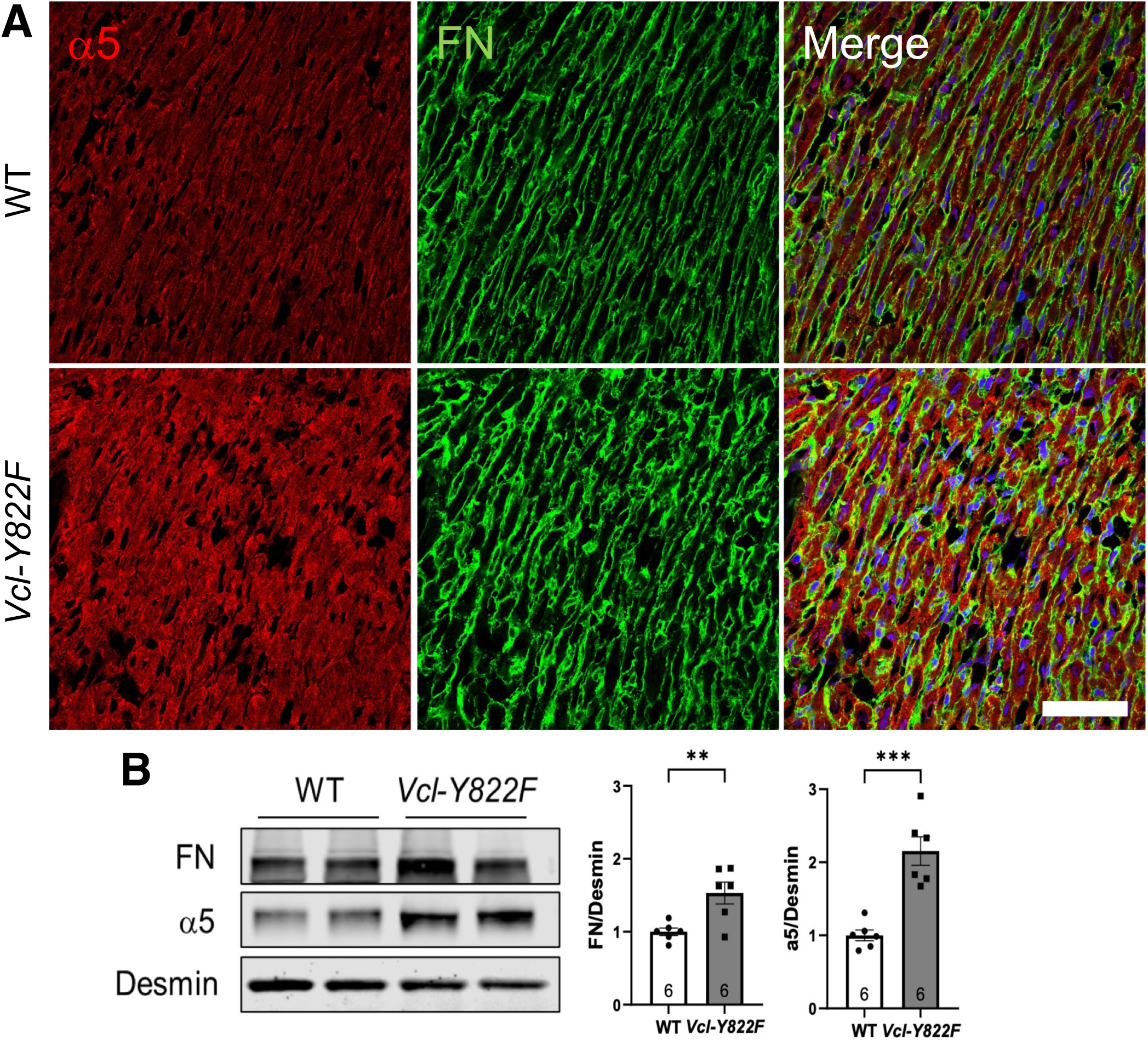
Increased cell-ECM adhesions in Vcl Y822F hearts. (A) Representative immunofluorescence images of P7 heart sections co-stained with α5 integrin (red) and fibronectin (FN, green). Scale bar: 50 μm. (B) Hearts were isolated from P7 mice (n=6/genotype) and Western blot analysis was performed on DOC insoluble heart lysates. Desmin was used as a loading control. **, p<0.01; ***, p<0.001 by two-tailed unpaired Student’s t-test. Error bars represent SEM.

### pY822 regulates VCL recruitment to cell-cell contacts in cultured neonatal CMs

A hallmark of CM maturation in the developing heart is the transition from an immature, unorganized, and rounded morphology to a mature, well-organized, and rod-shaped morphology. This maturation coincides with significant changes in the cardiac microenvironment, notably a transition from soft FN in the embryonic heart to stiff collagen I (COL) in mature tissue. Our recent studies have shown that VCL enrichment at adhesion complexes is ECM-dependent. Notably, we observe a significant enrichment of VCL at AJs when CMs are cultured on PDMS substrates coated with COL versus FN **(Fig. 5A)**. To determine if pY822 regulates ECM-dependent VCL enrichment at adhesion complexes, we isolated CMs from homozygous mutant *Vcl Y822F* neonates (isogenic C57BL/6J) and CMs isolated from C57BL/6J mice (referred to as wild type, WT). We plated CMs on FN and COL-coated PDMS substrates and stained them for VCL and plakoglobin to assess recruitment to AJs. We detected similar levels of VCL recruitment to AJs in WT and *Vcl Y822F* CMs on FN **(Fig. 5B)**. Strikingly, VCL AJ enrichment on COL was lost in Y822F CMs and was lower than observed on FN **(Fig. 5B)**. Given that COL is the primary matrix protein in P28 hearts, the reduction in VCL recruitment to AJs is similar to the loss of VCL at the ICD in *Vcl Y822F* mouse hearts (**Fig. 3**). Furthermore, it is consistent with our hypothesis that pY822 is essential for regulating VCL functions in CM AJ organization and maturation.

**Figure 5.**
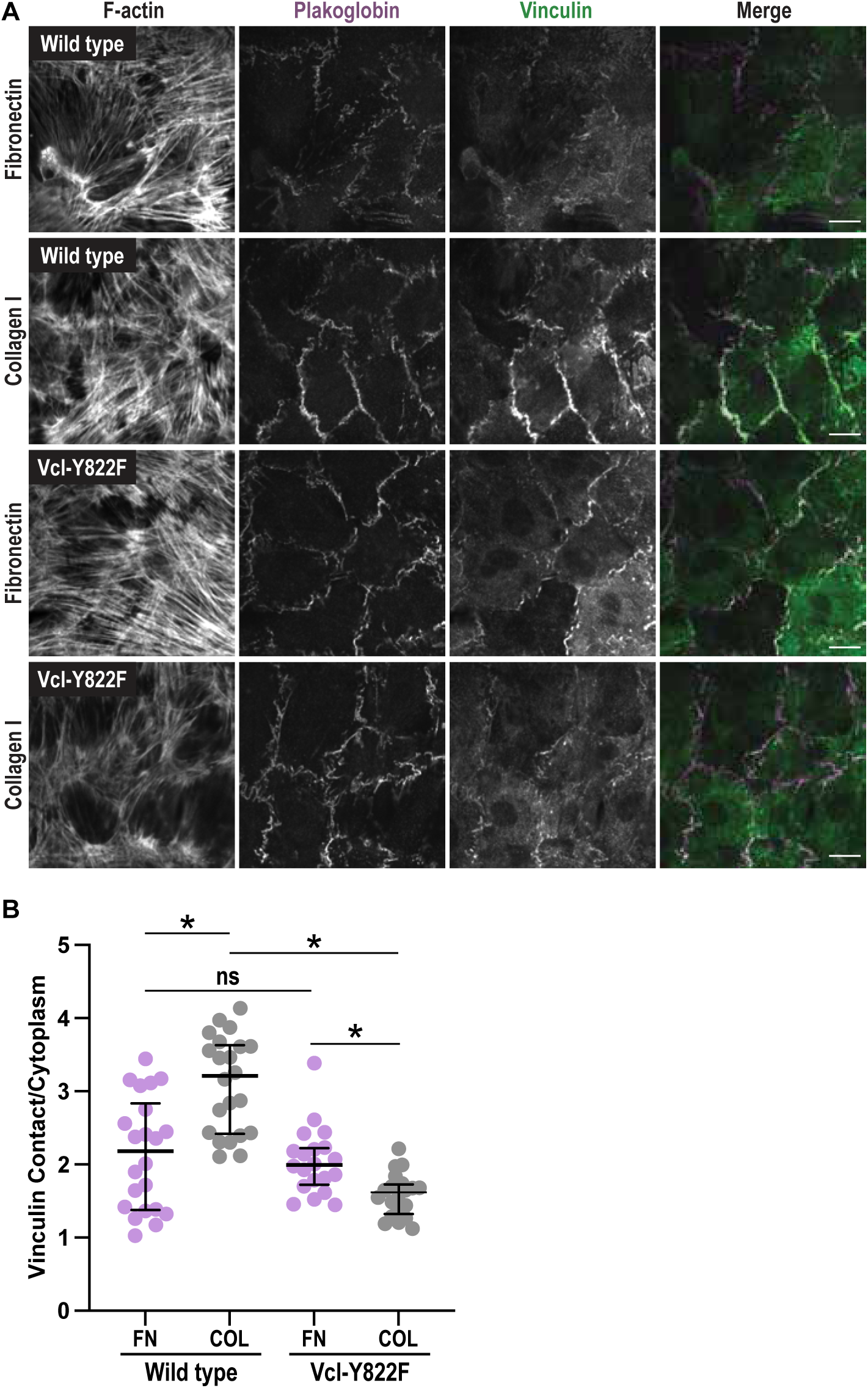
Y822 phosphorylation regulates vinculin recruitment to cell-cell contacts in cultured CMs. (A) Wild-type or *Vcl Y822F* CMs plated on fibronectin (FN) or collagen I (COL) coated PDMS, fixed 72 hrs post-plating, and stained for F-actin, plakoglobin, and vinculin. Scale bar 10 µm. (B). VCL contact to cytoplasmic signal ratio. One-way ANOVA (C) used to measure significance.

### VCL Y822F dynamics are similar to VCL WT

The reduced VCL recruitment to AJs in Y822F CMs observed *in vivo* and *in cellulo* led us to question whether the Y822F mutation impacted VCL localization and dynamics at CM FAs and AJs. To test this, mEmerald(Em)-tagged VCL and VCL Y822F were individually transfected into cultured neonatal WT CMs. As expected, Em-VCL localized to cardiomyocyte FAs and AJs **(Fig. S3)**. Em-VCL Y822F also localized to both FAs and AJs **(Fig. S3)**. Protein dynamics were measured using fluorescent recovery after photobleaching (FRAP) in dense cultures plated for 48-72 hours (**30**). Fluorescence recovery at FAs and AJs over ten minutes was quantified, plotted, and fit to a double exponential curve **(Fig. 6A-D)**. The mobile fractions of VCL and VCL Y822F at FAs (45% and 50%, respectively) and AJs (42% and 49%, respectively) were similar **(Fig. 6E)**. We then assessed the recovery rates for both the fast and slow pools of VCL and VCL Y822F. The fast pool recovery halftimes (4.9 – 21.9 seconds, secs) likely reflect an unbound, cytosolic population of VCL protein near cell junctions. Alternatively, the pool could be caused by photoswitching (**35**). Importantly, the fast pool for both VCL and VCL Y822F at both AJs and FAs was relatively small (13.6 – 18.8%, **Fig. 6E**), and the slow pool represents the dynamics of most of the junctional population. Like the mobile fraction, the recovery halftimes of VCL and VCL Y822F were similar at AJs (279.6 and 231.0 secs, respectively) and FAs (206.8 and 238.4 secs, respectively) **(Fig. 6E)**. We conclude that VCL dynamics at both junction types are 1) similar despite distinct binding partners and 2) the Y822F mutation does not affect protein dynamics at stable junctions. Notably, the VCL and VCL Y822F mobile fractions are larger, and recovery rates slower, in CMs than those reported in non-muscle cells (36–38), indicating that VCL is more stably associated with CM AJs and FAs. Given that load increases VCL binding to F-actin and VCL stability at junctions (16, 39), contraction forces likely regulate VCL dynamics at CM junctions.

**Figure 6.**
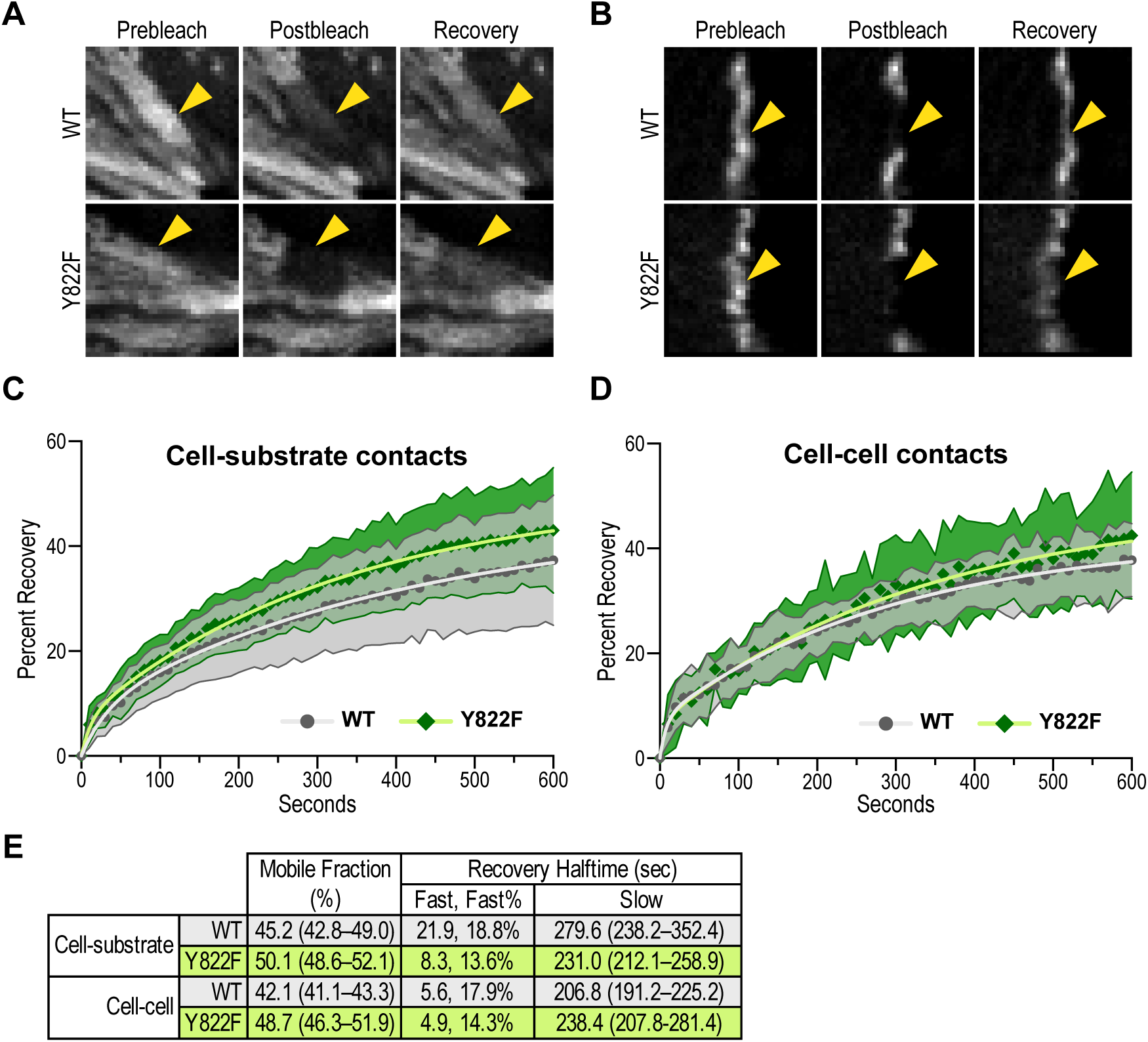
VCL Y822F dynamics are similar to VCL WT. A,B. Images from FRAP experiments showing pre-bleach, post-bleach, and recovery after 10 minutes in cardiomyocytes expressing Emerald-tagged VCL WT or VCL Y822F at cell-substrate contacts (A) or cell-cell contacts (B). C, D. Plots of mean +/− standard deviation of FRAP recovery fraction over time for Emerald-VCL WT (WT, dark grey circles) and Emerald-vinculin Y822F (green diamonds). Recovery plots are shown for cell-substrate (C) and cell-cell (D) contacts. The data were fit to a double exponential curve (WT, light grey line; Y822F, light green line). The number of experimental replicates/FRAP regions quantified: cell substrate (C) WT 2/32, Y822F 2/32; cell-cell (D) WT 3/17, Y822F 4/13. E. Summary of the mobile fraction (percentage), recovery halftimes (fast and slow pools), and the fast pool percentage of total recovered protein. The 95% confidence interval for mobile fraction percentage and slow pool recovery halftime is shown.

### *Vcl Y822F* mutant mice exhibit cardiac dysfunction

Next, we examined the cardiac phenotype in adult *Vcl Y822F* mutant mice. We did not detect any morphological or histological abnormalities in the Vcl Y822F hearts at 28 weeks of age **(Fig. 7A, B)**. Staining with Masson’s Trichrome revealed no apparent fibrosis in the *Vcl* mutant hearts **(Fig. S4)**. Moreover, we found no difference in heart weight/body weight or heart weight/tibia length between the WT and *Vcl Y822F* mutant mice at 40 weeks of age **(Fig. 7C, D)**. To assess cardiac function, we performed echocardiography on *Vcl* mutant mice at 10 and 28 weeks of age **(Fig. 7E-H**, **Table I)**. No difference in LV ejection fraction (EF) or fraction shortening (FS) was observed in the *Vcl Y822F* mice at 10 weeks of age. However, the *Vcl* mutant mice exhibited a reduction in LV function with age **(Fig. 7E-H**, **Table I)**. These data provide the first *in vivo* evidence that VCL Y822 phosphorylation regulates cardiac function in older animals.

**Figure 7.**
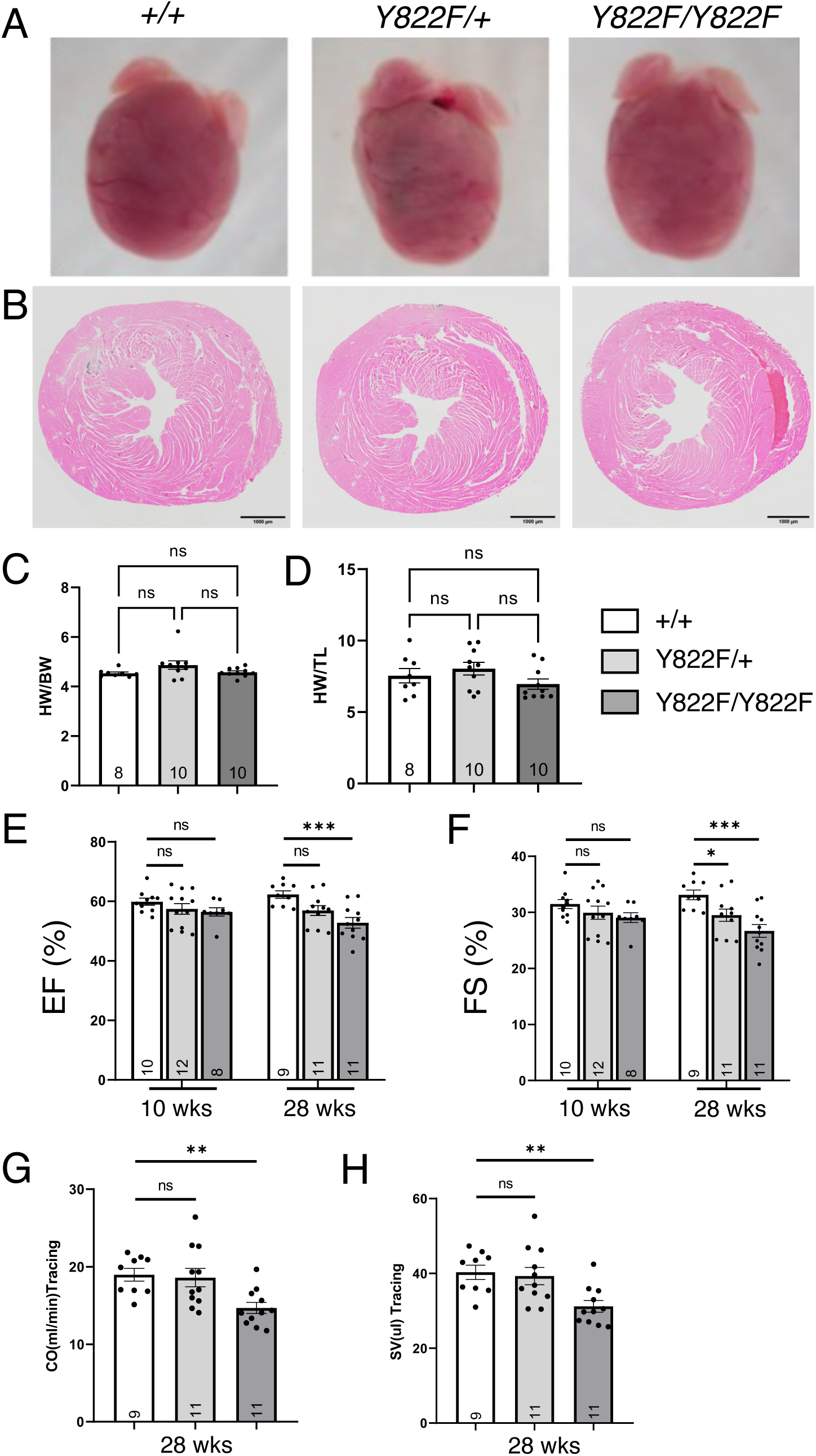
Histological and cardiac function analysis of *Vcl Y822F* hearts. (A) Representative wholemount images of hearts from *Vcl +/+*, *Vcl Y822F/+,* and *Vcl Y822F/Y822F* mice at 40 wks of age. (B) Representative H&E-stained heart sections from 28-week-old mice. Quantification of heart weight/body weight ratio (C) and heart weight/tibia length ratio (D) at 40 weeks of age. Serial echocardiographic analyses were performed on mice from 10 - 28 wks of age. Comparison of (E) ejection fraction (EF) and (F) fraction shortening (FS) at 10 and 28 wks of age, and (G) cardiac output (CO) and (H) stroke volume (SV) at 28 weeks of age is shown.

**Table I.** Echocardiographic indices of cardiac size and function.

Echocardiographic indices of cardiac size and function
|  | WT | Y822 F/+ | Y822 F/F |
| --- | --- | --- | --- |
| n | 9 | 11 | 11 |
| Gender(M/F) | 4/5 | 7/4 | 4/7 |
| Age(mo) | 7 | 7 | 7 |
| Heart rate(bpm) | 471.44±3.95 | 472.27±4.05 | 470.82±2.75 |
| LVAW;d(mm) | 0.95±0.02 | 1.00±0.02 | 0.98±0.03 |
| LVAW;s(mm) | 1.10±0.03 | 1.15±0.02 | 1.14±0.02 |
| LVID;d(mm) | 3.80±0.11 | 3.96±0.10 | 3.75±0.07 |
| LVID;s(mm) | 2.58±0.11 | 2.82±0.09 | 2.68±0.08 |
| LVPW;d(mm) | 0.76±0.02 | 0.78±0.03 | 0.76±0.01 |
| LVPW;s(mm) | 1.14±0.02 | 1.15±0.02 | 1.10±0.03 |
| CO (LV Trace) (mL/min) | 18.96±0.82 | 18.60±1.18 | 14.68±0.72** |
| Diameter;d (LV Trace) (mm) | 3.87±0.10 | 3.97±0.10 | 3.72±0.06 |
| Diameter;s (LV Trace) (mm) | 2.59±0.09 | 2.80±0.09 | 2.73±0.06 |
| EF (LV Trace) (%) | 62.30±1.27 | 56.91±1.65 | 52.80±1.79*** |
| FS (LV Trace) (%) | 33.12±0.86 | 29.51±1.09* | 26.70±1.13*** |
| LV Mass(mg) | 119.38±5.81 | 134.13±6.13 | 119.61±3.78 |
| LV Mass (Corrected) (mg) | 95.50±4.65 | 107.30±4.90 | 95.69±3.03 |
| LV Vol;d(μL) | 62.79±4.42 | 69.12±4.17 | 60.48±2.70 |
| LV Vol;s(μL) | 24.77±2.61 | 30.58±2.37 | 26.88±2.00 |
| SV (LV Trace) (μL) | 40.29±1.88 | 39.28±2.31 | 31.19±1.54** |
| V;d (LV Trace) (μL) | 65.17±3.92 | 69.32±4.03 | 59.17±2.20 |
| V;s (LV Trace) (μL) | 24.88±2.20 | 30.04±2.27 | 27.99±1.50 |

LVID, left ventricular internal dimension; LV vol, left ventricular volume; EF, ejection fraction; FS, fraction shortening; LVAW, left ventricular anterior wall; LVPW, left ventricular posterior wall; d, diastole; s, systole; CO, cardiac output; SV, stroke volume.
LV Trace: using Trace method. Values are represented as Mean±SEM.
One-way ANOVA with Dunnett's multiple comparison, vs WT group,
\* $p < 0.05$ , \*\* $p < 0.01$ , \*\*\* $p < 0.001$ .

## Discussion

The remodeling of myocardial cell-ECM and cell-cell adhesions is essential for mechanical coordination and tissue organization during cardiac development. In addition, disruptions in cell-ECM and cell-cell adhesion homeostasis are linked to heart disease and a hallmark of aging. Vinculin phosphorylation has emerged as a critical posttranslational regulator of adhesive activity at FAs and AJs in cultured cells (6, 10). However, the *in vivo* role of VCL phosphorylation in physiology and disease is poorly defined. The postnatal heart offers a unique model for investigating junction remodeling during CM maturation. Myocardial AJs and FAs reorganize as CMs adapt to increased cardiac demand and growth, and this reorganization requires coordinated and compensatory changes in their molecular organization and mechanical properties. Here, for the first time, we provide evidence that phosphorylation is critical for regulating VCL function in CM junction organization both *in vivo* and *in cellulo*.

We found that pY822 VCL expression was highest in the developing and perinatal heart but decreased significantly after two weeks of age and remained low in the adult heart. This expression pattern correlates with the profound rearrangement of cell adhesions during CM maturation. This reorganization culminates with AJs localized to the longitudinal termini and FAs restricted to the lateral membrane of the differentiated CM. We speculate that pY822 VCL regulates the organization of the dynamic, transitory adhesion complexes that promote junction maturation in CMs.

The expression and cellular distribution of pY822 VCL depends on multiple factors: cell type (e.g., nonmuscle vs. muscle cell), maturation state of the cadherin adhesion complex, and cytoskeletal tension. In the developing and neonatal heart, pY822 VCL was predominantly nonjunctional, displaying a punctate and diffuse staining pattern. In contrast, in confluent cultures of mammary epithelial cells, pY822 VCL largely co-localized with E-cadherin at cell-cell contacts (11). However, in other *in vitro* systems, pY822 VCL was largely punctate and nonjunctional. Notably, pharmacological enhancement of barrier function in cultured endothelial cells increased pY822 VCL at cell-cell contacts (40). Similarly, pY822 VCL was recruited to cell-cell contacts in cultured corneal epithelial cells after scratch wounding (41). Since these cell manipulations induce profound changes in cytoskeleton organization and cell tension (42), we speculate that increased actomyosin tension promotes VCL Y822 phosphorylation and concomitant recruitment to cell-cell contacts. Consistent with this, we previously reported that activation of nonmuscle myosin contractility promoted pY822 VCL expression and recruitment to neonatal CM cell-cell contacts (43). Together, these results suggest that cytoskeletal force regulates VCL Y822 phosphorylation. These data also indicate the existence of three distinct pools of VCL in cells: AJs, FAs, and a less appreciated nonjunctional pool. The function of the nonjunctional pool remains unclear, though recent work suggests that it is under load (16). Future work is expected to determine how VCL cellular distribution is regulated and if or how changes in distribution impact cell adhesion and organization.

To investigate the role of pY822 *in vivo*, we engineered a phosphodeficient *Vcl* mouse (Y822F) using CRISPR/Cas9 genome editing. Loss of Y822 phosphorylation caused a reduction in VCL at myocardial cell-cell contacts and disrupted AJ maturation in the postnatal heart. Cell-cell adhesions were also perturbed in cultured Y822F neonatal CMs isolated from the *Vcl* mutant mice. Interestingly, while the Y822F mutation reduced VCL recruitment to AJs, FRAP analysis showed that the Y822F mutation did not affect VCL dynamics at either cell-ECM or cell-cell adhesions. While these findings appear contradictory, they likely reflect that only a small fraction of the total pool of VCL is phosphorylated, even when pY822 levels are at their highest. Thus, Y822F (unphosphorylated VCL) represents the “dominant” species of VCL in the cell, and therefore, its dynamics should mimic wild-type VCL since most wild-type protein is unphosphorylated at Y822.

Studies of cultured CMs have shown that integrins reinforce myocardial cell-cell adhesions (44). Integrins organize adjacent to nascent cell-cell adhesions when neonatal CMs are grown on micropatterned FN substrates, suggesting that the two adhesion structures cooperate to generate the ICD (44). Here, we show that myocardial cell-ECM adhesions were enhanced in *Vcl Y822F* neonatal hearts. Specifically, α5/β1 integrin and FN expression were increased. Given the defects in AJ organization, FA strengthening in *Vcl Y822F* hearts suggests a compensatory mechanism. Note that loss of N-cadherin in adult myocardium leads to upregulation of β1 integrin (45), consistent with such a compensatory response. Collectively, these data highlight the functional interplay between cadherin and integrin adhesions and underscore the importance of VCL in coordinating adhesion organization and maintaining mechanical homeostasis in the heart.

Multiple phosphorylation sites have been implicated in VCL function at FAs and AJs. Src kinase-mediated phosphorylation of VCL (Y100/Y1065) is important for platelet activation and spreading (32). In airway smooth muscle tissue, VCL phosphorylation at tyrosine residue 1065 results in an “open”, active conformation that promotes actin polymerization in response to contractile stimulation (46). Conversely, expression of non-phosphorylatable VCL Y1065F decreased actin polymerization and contractile force in smooth muscle cells (46). During collective cell migration of MDCK cells, phosphorylation at S1033 biases VCL toward a closed, unloaded state at AJs and within the cytoplasm (16). A phosphoproteomic study recently identified a new VCL phosphorylation site, serine 721 (pS721), associated with a porcine atherosclerosis model (47). Notably, VCL is upregulated in atherosclerosis, detectable in patient serum, and differentially expressed in unstable versus stable atherosclerotic plaques (47–49). S721 is phosphorylated in endothelial cells (ECs) undergoing disturbed shear flow. In response to disturbed flow, pS721 inactivates VCL by maintaining it in a closed conformation, leading to disrupted VE-cadherin adhesion, increased endothelial permeability, and atherogenesis (47). EC-specific expression of a non-phosphorylatable *Vcl S721A* in the apolipoprotein E-deficient (ApoE -/-) mouse model suppressed plaque formation (47). Hence, VCL phosphorylation may be a widely used mechanism to respond to changes in mechanical forces in cardiovascular disease.

Age modifies heart structure and function, even without overt cardiovascular disease. Notably, VCL is overexpressed in healthy aged myocardium from flies to non-human primates (50). In the hearts of aged rats and rhesus monkeys, VCL accumulated at the AJ, similar to humans with heart failure (51). Engler and colleagues showed that VCL is a unique molecular regulator of cardiac function during aging in flies. Cardiac-specific overexpression of VCL induced cytoskeletal and cellular remodeling that correlated with increased cardiac contractility, energy metabolism, organismal fitness, and extended *Drosophila* life span. Thus, VCL-mediated cytoskeletal remodeling is cardioprotective in the aged heart (50, 52). It will be interesting to determine if posttranslational modification of VCL occurs with age and whether cardiac dysfunction is exacerbated in 2-year-old *Vcl Y822F* mice.

In conclusion, our study provides *in vivo* evidence for VCL post-translational modification as a critical coordinator of AJ and FA remodeling in CMs. Loss of Y822 phosphorylation results in aberrant AJ maturation and increased α5-FN expression, consistent with the idea that CMs rebalance their cell-cell and cell-ECM adhesions to restore mechanical homeostasis in the heart. There are many unanswered questions about the function of pY822 VCL in the cell. What is the role of the nonjunctional pool in myocardial junction maturation? What are the binding partners of pY822 VCL, and how do they regulate its activity? Also, it will be interesting to determine whether pY822 VCL is cardioprotective following injury and, if so, whether modifying VCL phosphorylation influences the pathophysiology of heart disease. Future experiments will be necessary to address these and other questions.

## Materials and Methods

### Mouse model generation by CRISPR-mediated gene editing, genotyping, and colony maintenance

CRISPR-targeting reagents were injected into the cytoplasm of the C57BL/6J mouse zygotes by the Brown University Mouse Transgenic and Gene-Targeting Facility. The injection mix contained 100 ng μl^−l^ SpCas9 nuclease (Cat. No. 1081060, Integrated DNA Technologies (IDT)), 150 ng μl^−l^ the annealed guide RNA (crRNA:tracrRNA, IDT), and 250 ng μl^−l^ single-stranded DNA oligonucleotide carrying the targeted mutation *Vcl^Y822F^*(TAT >TtT) (IDT) and a silent mutation to disrupt the targeting site PAM domain (Supplemental Table II). gRNA sequence close to the mutation site and with minimum potential off-target cleavage activity was selected (Supplemental Table III). Mosaic *Vcl^Y822F/+^* founder mice were bred with wild-type C57BL/6J mice for four generations to diminish potential off-target mutations. Two independent *Vcl^Y822F/+^* founder lines were intercrossed to generate homozygous *Vcl^Y822F/Y822F^* mice. Mice were genotyped by *Vcl* gene locus-specific PCR. The *Y822F* mutant allele resulted in the loss of an EcoRV site at the *Vcl* locus (GATATC >GAT**T**TC. Following EcoRV digestion of the PCR product (514 bp), the *Vcl^+/+^*genotype corresponds to two DNA bands (273 and 241 bps), *Vcl^Y822F/+^*has the full-length DNA band (514 bps) plus the two bands, and *Vcl^Y822F/V822F^*has only the undigested PCR product. To ensure inclusion of both sexes for these studies, the sex of neonatal pups was confirmed by the presence or absence of the Y-chromosome via PCR and/or observation of the anogenital distance. All animal studies were performed in accordance with the guidelines of the IACUC of Rhode Island Hospital.

### Echocardiographic and hemodynamic measurements

Transthoracic two-dimensional echocardiography was performed on *Vcl^+/+^*, *Vcl^Y822F/+^*, and *Vcl^Y822F/Y822F^* mice with a Vevo 2100 ultrasound system equipped with a 30-MHz transducer (FUJIFILM VisualSonics Inc., Toronto, ON, Canada). Heart rate (HR), ejection fraction (EF%), fractional shortening (FS%), left ventricular internal dimensions (LVID), left ventricular volume (LV vol), LV posterior wall thickness (LVPW), LV anterior wall thickness (LVPW), were measured from M-mode images at the plane bisecting the papillary muscles. The researcher was blinded to the genotype of the mice.

### Histological analysis

Mice hearts were isolated and washed in PBS. The hearts were then fixed in fresh 4% paraformaldehyde at 4°C overnight, dehydrated, and embedded in paraffin. Longitudinal 5 μm deparaffinized sections were stained with Hematoxylin and Eosin (H&E) or Masson Trichrome, mounted and imaged with an Olympus BX43 microscope system. Images were processed with cellSens™ Imaging Software.

### Immunofluorescence

Heart tissues were harvested and embedded in OCT (Sakura Finetek). OCT sections (6 μm) were fixed in 4% paraformaldehyde (PFA) for 10min and then washed with PBS 3 times. The heart sections were then permeabilized in 0.1% Triton X-100 (in PBS) for 10min at room temperature. Sections were blocked in T-BSA (5% BSA, 0.01% Triton X-100 in PBS) for 1 hour at room temperature and incubated overnight at 4°C with primary antibodies: N-cadherin (1:400, 610921, BD Bioscience), α5 integrin (1:200, 553319, BD bioscience), fibronectin (1:100, PA1-23693, Thermofisher), pY822 VCL (pVCL) (1:200, ab61071/ab200825, Abcam; or 1:200, 44-1080G, Invitrogen), VCL (1:100, V4505, Sigma), VCL (1:100, V9131, Sigma), α-actinin (1:200, A7732, Sigma), β-catenin(1:200, C2206, Sigma). After washing in PBS, sections were incubated with secondary antibody (Alexa Fluor (A) 488 goat anti-rabbit or A488 goat anti-mouse IgG, A555 goat anti-mouse IgG or donkey anti-rabbit IgG, A586 donkey anti-rat IgG, Life Technologies) for 1 hour at room temperature. Finally, the tissue sections were washed in PBS and mounted with ProLong Gold Antifade Reagent containing DAPI (Life Technologies). Images were acquired using a Nikon A1R Confocal Microscope System. Images were processed and analyzed using Nikon NIS-Elements software and FIJI ImageJ 1.53 software.

### Western blot analysis

The harvested mice heart tissues were homogenized in a modified RIPA buffer (50mM Tris-HCl pH 7.5, 150mM NaCl, 1mM EDTA pH 8.0, 1% NP-40, 0.5% Na deoxycholate, 0.1% SDS) containing protease inhibitor and phosphatase inhibitor cocktails II and III (Roche Diagnostics). After rotating at 4°C for 2 hours, the mixtures were centrifuged at 13,000g for 15 min at 4°C. Primary antibodies used were: α5 integrin (1:200, sc-166681, Santa Cruz), pY822 VCL (pVCL) (1:1000, ab61071/ab200825, Abcam; or 1:1000, 44-1080G, Invitrogen), VCL (1:1000, V4505, Sigma), N-cadherin (1:1000, 610921, BD Bioscience), GAPDH (1:3000, 6c5, RDI), afadin (1:2000, A0349, Sigma), fibronectin (1:3000, PA1-23693, Thermo Fisher), αE-catenin(1:50, 711200, Invitrogen), αT-catenin(1:100, 13974-1-AP, Proteintech), β-catenin(1:200, C2206, Sigma), pY100 VCL (1:1000, 44-2078G, Invitrogen), pY1065 VCL (1:1000, 44-1074G, Invitrogen). For normalization of signals, blots were also analyzed with anti-GAPDH antibody followed by IRDye 680 or IRDye 800CW conjugated secondary antibody (1:15000, LI-COR, Lincoln, NE). anti-mouse-HRP or anti-rabbit-HRP secondary antibody (1:3000, Bio-Rad) were also used and developed by SuperSignal West Pico PLUS Chemiluminescent Substrate (Thermo Scientific, Rockford, IL). Membranes were imaged using the Odyssey Infrared Imaging System (LI-COR) or ChemiDoc Imaging System (Bio-Rad, Hercules, CA). Quantification was performed with FIJI ImageJ 1.53 software.

### Preparation of Deoxycholate (DOC) soluble and insoluble fractions

Deoxycholate (DOC) soluble and insoluble fractions of harvested heart tissues were prepared as previously reported (53). Briefly, fresh heart tissues were homogenized in DOC lysis buffer (Tris-HCl (pH 8.8) 20mM, EDTA 2mM, PMSF 2mM, Iodoacetic acid 2mM, N-ethylmaleimide 2mM, DOC 2%) with protease inhibitors, and the lysates were centrifuged at 13,000g 4°C for 15 min. The supernatants were saved as the DOC soluble fraction. The pellets were re-suspended in SDS solubilization buffer (Tris-HCl (pH 8.8) 20mM, EDTA 2mM, PMSF 2mM, Iodoacetic acid 2mM, N-ethylmaleimide 2mM, SDS 1%) with protease inhibitors. The protein concentration of DOC soluble fractions was measured by BCA assay. DOC insoluble sample volume for SDS-PAGE is based on protein concentration in the corresponding DOC soluble fraction.

### Plasmids

mEmerald-Vinculin-23 was a gift from Michael Davidson (Addgene plasmid # 54302). mEmerald-vinculin Y822F was made by site-directed mutagenesis.

### Cardiomyocyte Isolation and Culture

All animal work was approved by the University of Pittsburgh Division of Laboratory Animal Resources. Outbred Swiss Webster mice were used to generate wild-type CMs for transfection and FRAP experiments. Inbred B6 mice were used to create wild-type control CMs for immunostaining experiments. *Vcl^Y822F/V822F^* mice were used to generate Y822F homozygous mutant cardiomyocytes.

MatTek dishes (35 mm dish with 10 mm microwell) were coated with rat tail Type I collagen (Millipore) diluted to 0.5 μg/μl in PBS for 1 hour at room temperature. The dishes were dried and treated with UV radiation for 1 hour, after which they were washed with PBS, dried, and stored at room temperature in the dark.

PDMS substrates were prepared by mixing twenty parts of silicone elastomer base with one part of curing agent (SYLGARD, Dow Corning, USA) and stirred. The mixture was degassed for 20 minutes in a vacuum chamber to remove air bubbles. 22 x 22 mm glass coverslips (Corning) were cleaned by sequentially rinsing in ethanol and water, then flaming to sterilize. Sterilized coverslips were spin-coated with PDMS (Laurell Technologies Corporation Model WS-400B-6NPP/LITE/8K) at 5000 rpm for 1 min and baked at 60 °C for 2 hours. PDMS-coated coverslips were washed in ethanol for 4 hours, followed by water overnight. The next day, PDMS-coated coverslips were dried with N2 gas flow and then activated with oxygen plasma (Harrick Plasma Cleaner, PDC-001) for 3 minutes. Activated coverslips were treated with 2% (3-Aminopropyl) triethoxysilane (APTES) dissolved in 95% ethanol for 20 min at 60 °C. Coverslips were washed in ethanol ten times, followed by distilled water ten times. Immediately after surface activation and treatment, Type I collagen (0.5 μg/μl, Millipore) or fibronectin (0.025 µg/µl, Santa Cruz) was added, and coverslips were incubated for 1 hour at room temperature. ECM-coated coverslips were stored at 4°C until use.

Neonatal mouse cardiomyocytes were isolated as described (54). Briefly, mouse pups were sacrificed at P1, and the hearts were removed, cleaned, minced, and digested overnight at 4°C in 20 mM BDM (2,3-Butanedione monoxime) and 0.0125% trypsin in HBSS. The following day, heart tissue was digested in 15 mg/mL Collagenase/Dispase (Roche) in Leibovitz media with 20 mM BDM to create a single-cell suspension. Cells were pre-plated for 1.5 hours in plating media (65% high glucose DMEM, 19% M-199, 10% horse serum, 5% FBS, and 1% penicillin-streptomycin) to remove fibroblasts and endothelial cells. Cardiomyocytes were plated on MatTek dishes (1.5×10^5^) or PDMS substrates in plating media. 16 hours post-plating, the plating media was exchanged for maintenance media (78% high glucose DMEM, 17% M-199, 4% horse serum, 1% penicillin-streptomycin, 1 μM AraC, and 1 μM isoproterenol).

### Immunofluorescence

CMs were processed for immunofluorescence as follows: cells were fixed in warmed (37 °C) 4% EM grade paraformaldehyde in PBS (Ca^2+^, Mg^2+^) and 0.12 M sucrose for 10 minutes and washed twice with PBS (Ca^2+^, Mg^2+^). Cells were permeabilized with 0.2% Triton X-100 in PBS (Ca^2+^, Mg^2+^) for 5 minutes and washed twice with PBS (Ca^2+^, Mg^2+^). Cells were blocked in 10% BSA (Sigma) in PBS (Ca^2+^, Mg^2+^) for 1 hour at room temperature. Samples were incubated with primary antibodies in PBS (Ca^2+^, Mg^2+^) + 1% BSA for 1 hour at room temperature, washed 2X in PBS (Ca^2+^, Mg^2+^), incubated with secondary antibodies in PBS (Ca^2+^, Mg^2+^) + 1% for 1 hour at room temperature, washed 2X in PBS (Ca^2+^, Mg^2+^), and then mounted in Prolong Glass (Thermo Fisher Scientific). All samples were cured at least 24 hours before imaging.

### Antibodies for cell staining

Primary antibodies used for immunostaining were: anti-αE-catenin (1:100; Enzo Life Sciences ALX-804-101-C100), anti-Plakoglobin (1:100; Cell Signaling 2309), anti-Vinculin (1:800; Sigma Aldrich V9131), anti-N-cadherin (1:250; Invitrogen 99-3900), Secondary antibodies used were goat anti-mouse or anti-rabbit IgG conjugated to A488, A568 or A647 (1:250; Invitrogen). F-actin was visualized using an Alexa Fluor dye conjugated to phalloidin (1:100, ThermoFisher Scientific).

### Confocal Microscopy

Cells were imaged with a 100X 1.49 NA objective on a Nikon Eclipse Ti inverted microscope outfitted with a Prairie swept field confocal system, Agilent monolithic laser launch, and Andor iXon3 camera using NIS-Elements (Nikon) imaging software. Maximum projections of 2-3 µm of the image stacks were created for presentation. Expression and staining levels were adjusted for presentation purposes in Photoshop (Adobe). All levels were corrected the same across each figure.

### FRAP Experiments

FRAP experiments were performed on a Nikon swept field confocal microscope (described above) outfitted with a Tokai Hit cell incubator and Bruker miniscanner. Actively contracting cells were maintained at 37°C in a humidified, 5% CO_2_ environment. User-defined regions along cell-cell contacts were bleached with a 405 laser, and recovery images were collected every 10 seconds for 10 minutes. FRAP data was quantified in ImageJ (NIH), and average recovery plots were measured in Excel (Microsoft). All recovery plots represent data from at least two independent transfections of unique CM preps. The data were fit to a double-exponential curve to determine the mobile fraction and half-time of recovery in Prism (GraphPad).

### Image Analysis

To quantify vinculin recruitment to CM cell-cell contacts, a maximum intensity projection of 6 z-stack slices containing cell-cell contacts was generated in ImageJ. IsoJ Dark thresholding was then preformed on the projected image to create a mask of the plakoglobin channel to define the region of analysis. Vinculin signal intensity was then measured within the masked region. Next, three random measurements of vinculin signal intensity in the cell cytoplasm were collected and averaged. Finally, the ratio of vinculin intensity within the mask was divided by the average cytoplasmic signal to normalize between samples and calculate the contact/cytoplasmic ratio. Data were plotted with Prism software (GraphPad). A One-way ANOVA with multiple comparisons was performed to determine significance; p<0.05.

### Statistical analysis

Statistical differences were assessed by unpaired, two-tailed Student’s t-test or one-way ANOVA followed by Tukey’s multiple comparison of individual means. A *p*-value of < 0.05 was considered statistically significant.

## Supplemental figure legends

**Supplemental Figure 1. The protein levels of the N-cadherin/catenin complex do not change in the *Vcl Y822F* hearts.** Western blots and quantitative analysis of N-cadherin (Ncad), αE-catenin (αE-cat), αT-catenin (αT-cat), β-catenin (β-cat), Afadin, and VCL expression in heart lysates from WT (n=4) and *Vcl Y822F* (n=5) mice at P14 (A) and P28 (B).

**Supplemental Figure 2. Phosphorylation at VCL Y100 and Y1065 does not change in the *Vcl Y822F* hearts.** Western blots and quantitative analysis of pY100 and pY1065 expression in heart lysates from WT (n=4) and *Vcl Y822F* (n=4) P7 mice.

**Supplemental Figure 3. Emerald-VCL WT and Emerald-VCL Y882F localize to FAs and AJs in transfected CMs.** Emerald-tagged VCL WT (top row, green in merge) or VCL Y822F (bottom row, green in merge) were transfected into CMs plated on COL, fixed 48 hrs post-transfection, and stained for F-actin and plakoglobin (magenta in merge). Individual channels and Emerald/plakoglobin merge shown. Scale bar 10 µm.

**Supplemental Figure 4. No increase in fibrosis in *Vcl Y822F* hearts.** (A) Representative Masson’s Trichrome-stained heart sections from *Vcl +/+*, *Vcl Y822F/+,* and *Vcl Y822F/Y822F* female mice at 40 wks of age.

## Acknowledgement

We thank Christopher Liu for technical assistance, and Jinping Luo and the Brown University Transgenic Facility.

## References

1. DuFort CC, Paszek MJ, and Weaver VM. Balancing forces: architectural control of mechanotransduction. Nat Rev Mol Cell Biol. 2011;12(5):308–19.

2. Ingber DE. Mechanobiology and diseases of mechanotransduction. Ann Med. 2003;35(8):564–77.

3. Jaalouk DE, and Lammerding J. Mechanotransduction gone awry. Nat Rev Mol Cell Biol. 2009;10(1):63–73.

4. Lyon RC, Zanella F, Omens JH, and Sheikh F. Mechanotransduction in cardiac hypertrophy and failure. Circ Res. 2015;116(8):1462–76.

5. Atherton P, Stutchbury B, Jethwa D, and Ballestrem C. Mechanosensitive components of integrin adhesions: Role of vinculin. Exp Cell Res. 2016;343(1):21–7.

6. Bays JL, and DeMali KA. Vinculin in cell-cell and cell-matrix adhesions. Cell Mol Life Sci. 2017;74(16):2999–3009.

7. Leckband DE, and de Rooij J. Cadherin adhesion and mechanotransduction. Annu Rev Cell Dev Biol. 2014;30:291–315.

8. Grashoff C, Hoffman BD, Brenner MD, Zhou R, Parsons M, Yang MT, et al. Measuring mechanical tension across vinculin reveals regulation of focal adhesion dynamics. Nature. 2010;466(7303):263–6.

9. Leerberg JM, Gomez GA, Verma S, Moussa EJ, Wu SK, Priya R, et al. Tension-sensitive actin assembly supports contractility at the epithelial zonula adherens. Curr Biol. 2014;24(15):1689–99.

10. Goldmann WH. Role of vinculin in cellular mechanotransduction. Cell Biol Int. 2016;40(3):241–56.

11. Bays JL, Peng X, Tolbert CE, Guilluy C, Angell AE, Pan Y, et al. Vinculin phosphorylation differentially regulates mechanotransduction at cell-cell and cell-matrix adhesions. J Cell Biol. 2014;205(2):251–63.

12. Bertocchi C, Wang Y, Ravasio A, Hara Y, Wu Y, Sailov T, et al. Nanoscale architecture of cadherin-based cell adhesions. Nat Cell Biol. 2017;19(1):28–37.

13. Dumbauld DW, Lee TT, Singh A, Scrimgeour J, Gersbach CA, Zamir EA, et al. How vinculin regulates force transmission. Proc Natl Acad Sci U S A. 2013;110(24):9788–93.

14. LaCroix AS, Lynch AD, Berginski ME, and Hoffman BD. Tunable molecular tension sensors reveal extension-based control of vinculin loading. Elife. 2018;7.

15. Rothenberg KE, Scott DW, Christoforou N, and Hoffman BD. Vinculin Force-Sensitive Dynamics at Focal Adhesions Enable Effective Directed Cell Migration. Biophys J. 2018;114(7):1680–94.

16. Shoyer TC, Gates EM, Cabe JI, Urs AN, Conway DE, and Hoffman BD. Coupling during collective cell migration is controlled by a vinculin mechanochemical switch. Proc Natl Acad Sci U S A. 2023;120(50):e2316456120.

17. Zhou DW, Lee TT, Weng S, Fu J, and Garcia AJ. Effects of substrate stiffness and actomyosin contractility on coupling between force transmission and vinculin-paxillin recruitment at single focal adhesions. Mol Biol Cell. 2017;28(14):1901–11.

18. DeWane G, Cronin NM, Dawson LW, Heidema C, and DeMali KA. Vinculin Y822 is an important determinant of ligand binding. J Cell Sci. 2023;136(12).

19. Hirschy A, Schatzmann F, Ehler E, and Perriard JC. Establishment of cardiac cytoarchitecture in the developing mouse heart. Dev Biol. 2006;289(2):430–41.

20. Vite A, and Radice GL. N-cadherin/catenin complex as a master regulator of intercalated disc function. Cell Commun Adhes. 2014;21(3):169–79.

21. Mittal A, Pulina M, Hou SY, and Astrof S. Fibronectin and integrin alpha 5 play requisite roles in cardiac morphogenesis. Dev Biol. 2013;381(1):73–82.

22. Matsushita T, Oyamada M, Fujimoto K, Yasuda Y, Masuda S, Wada Y, et al. Remodeling of cell-cell and cell-extracellular matrix interactions at the border zone of rat myocardial infarcts. Circ Res. 1999;85(11):1046–55.

23. Sessions AO, and Engler AJ. Mechanical Regulation of Cardiac Aging in Model Systems. Circ Res. 2016;118(10):1553–62.

24. Xu W, Baribault H, and Adamson ED. Vinculin knockout results in heart and brain defects during embryonic development. Development. 1998;125(2):327–37.

25. Zemljic-Harpf A, Manso AM, and Ross RS. Vinculin and talin: focus on the myocardium. J Investig Med. 2009;57(8):849–55.

26. Zemljic-Harpf AE, Miller JC, Henderson SA, Wright AT, Manso AM, Elsherif L, et al. Cardiac-myocyte-specific excision of the vinculin gene disrupts cellular junctions, causing sudden death or dilated cardiomyopathy. Mol Cell Biol. 2007;27(21):7522–37.

27. Olson TM, Illenberger S, Kishimoto NY, Huttelmaier S, Keating MT, and Jockusch BM. Metavinculin mutations alter actin interaction in dilated cardiomyopathy. Circulation. 2002;105(4):431–7.

28. Vasile VC, Ommen SR, Edwards WD, and Ackerman MJ. A missense mutation in a ubiquitously expressed protein, vinculin, confers susceptibility to hypertrophic cardiomyopathy. Biochem Biophys Res Commun. 2006;345(3):998–1003.

29. Vasile VC, Will ML, Ommen SR, Edwards WD, Olson TM, and Ackerman MJ. Identification of a metavinculin missense mutation, R975W, associated with both hypertrophic and dilated cardiomyopathy. Mol Genet Metab. 2006;87(2):169–74.

30. Merkel CD, Li Y, Raza Q, Stolz DB, and Kwiatkowski AV. Vinculin anchors contractile actin to the cardiomyocyte adherens junction. Mol Biol Cell. 2019;30(21):2639–50.

31. Fukuda R, Gunawan F, Ramadass R, Beisaw A, Konzer A, Mullapudi ST, et al. Mechanical Forces Regulate Cardiomyocyte Myofilament Maturation via the VCL-SSH1-CFL Axis. Dev Cell. 2019;51(1):62–77 e5.

32. Zhang Z, Izaguirre G, Lin SY, Lee HY, Schaefer E, and Haimovich B. The phosphorylation of vinculin on tyrosine residues 100 and 1065, mediated by SRC kinases, affects cell spreading. Mol Biol Cell. 2004;15(9):4234–47.

33. Sechler JL, Takada Y, and Schwarzbauer JE. Altered rate of fibronectin matrix assembly by deletion of the first type III repeats. J Cell Biol. 1996;134(2):573–83.

34. Auernheimer V, Lautscham LA, Leidenberger M, Friedrich O, Kappes B, Fabry B, et al. Vinculin phosphorylation at residues Y100 and Y1065 is required for cellular force transmission. J Cell Sci. 2015;128(18):3435–43.

35. Mueller F, Morisaki T, Mazza D, and McNally JG. Minimizing the impact of photoswitching of fluorescent proteins on FRAP analysis. Biophysical journal. 2012;102(7):1656–65.

36. Lavelin I, Wolfenson H, Patla I, Henis YI, Medalia O, Volberg T, et al. Differential effect of actomyosin relaxation on the dynamic properties of focal adhesion proteins. PLoS One. 2013;8(9):e73549.

37. Wolfenson H, Bershadsky A, Henis YI, and Geiger B. Actomyosin-generated tension controls the molecular kinetics of focal adhesions. J Cell Sci. 2011;124(Pt 9):1425–32.

38. Carisey A, Tsang R, Greiner AM, Nijenhuis N, Heath N, Nazgiewicz A, et al. Vinculin regulates the recruitment and release of core focal adhesion proteins in a force-dependent manner. Curr Biol. 2013;23(4):271–81.

39. Huang DL, Bax NA, Buckley CD, Weis WI, and Dunn AR. Vinculin forms a directionally asymmetric catch bond with F-actin. Science. 2017;357(6352):703–6.

40. Birukova AA, Shah AS, Tian Y, Moldobaeva N, and Birukov KG. Dual role of vinculin in barrier-disruptive and barrier-enhancing endothelial cell responses. Cell Signal. 2016;28(6):541–51.

41. Onochie OE, Zollinger A, Rich CB, Smith M, and Trinkaus-Randall V. Epithelial cells exert differential traction stress in response to substrate stiffness. Exp Eye Res. 2019;181:25–37.

42. Abreu-Blanco MT, Watts JJ, Verboon JM, and Parkhurst SM. Cytoskeleton responses in wound repair. Cell Mol Life Sci. 2012;69(15):2469–83.

43. Li X, McLain C, Samuel MS, Olson MF, and Radice GL. Actomyosin-mediated cellular tension promotes Yap nuclear translocation and myocardial proliferation through alpha5 integrin signaling. Development. 2023;150(2).

44. McCain ML, Lee H, Aratyn-Schaus Y, Kleber AG, and Parker KK. Cooperative coupling of cell-matrix and cell-cell adhesions in cardiac muscle. Proc Natl Acad Sci U S A. 2012;109(25):9881–6.

45. Kostetskii I, Li J, Xiong Y, Zhou R, Ferrari VA, Patel VV, et al. Induced deletion of the N-cadherin gene in the heart leads to dissolution of the intercalated disc structure. Circ Res. 2005;96(3):346–54.

46. Huang Y, Day RN, and Gunst SJ. Vinculin phosphorylation at Tyr1065 regulates vinculin conformation and tension development in airway smooth muscle tissues. J Biol Chem. 2014;289(6):3677–88.

47. Shih YT, Wei SY, Chen JH, Wang WL, Wu HY, Wang MC, et al. Vinculin phosphorylation impairs vascular endothelial junctions promoting atherosclerosis. Eur Heart J. 2023;44(4):304–18.

48. Wu C, Li W, Li P, and Niu X. Identification of a hub gene VCL for atherosclerotic plaques and discovery of potential therapeutic targets by molecular docking. BMC Med Genomics. 2024;17(1):42.

49. Kristensen LP, Larsen MR, Mickley H, Saaby L, Diederichsen AC, Lambrechtsen J, et al. Plasma proteome profiling of atherosclerotic disease manifestations reveals elevated levels of the cytoskeletal protein vinculin. J Proteomics. 2014;101:141–53.

50. Kaushik G, Spenlehauer A, Sessions AO, Trujillo AS, Fuhrmann A, Fu Z, et al. Vinculin network-mediated cytoskeletal remodeling regulates contractile function in the aging heart. Sci Transl Med. 2015;7(292):292ra99.

51. Heling A, Zimmermann R, Kostin S, Maeno Y, Hein S, Devaux B, et al. Increased expression of cytoskeletal, linkage, and extracellular proteins in failing human myocardium. Circ Res. 2000;86(8):846–53.

52. Sessions AO, Min P, Cordes T, Weickert BJ, Divakaruni AS, Murphy AN, et al. Preserved cardiac function by vinculin enhances glucose oxidation and extends health-and life-span. APL Bioeng. 2018;2(3).

53. Wierzbicka-Patynowski I, Mao Y, and Schwarzbauer JE. Analysis of fibronectin matrix assembly. Curr Protoc Cell Biol. 2004;Chapter 10:Unit 10 2.

54. Ehler E, Moore-Morris T, and Lange S. Isolation and culture of neonatal mouse cardiomyocytes. J Vis Exp. 2013(79).

