## Supplemental Data for "Vinculin Y822 phosphorylation regulates cardiomyocyte adhesion dynamics and adherens junction maturation in the heart"

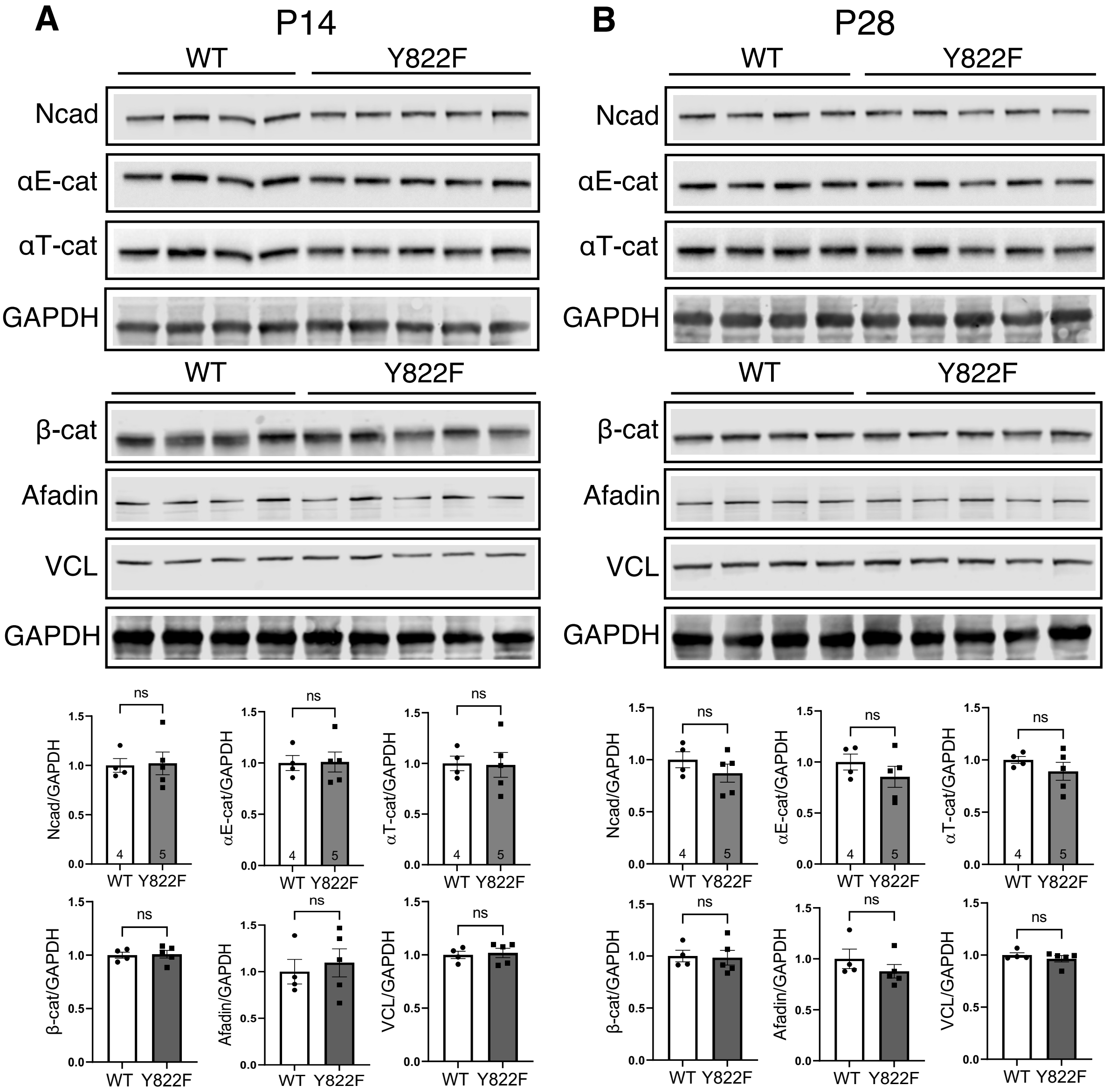


**Supplemental Figure 1.** Western blots and quantitative analysis of Ncad, αE-cat, αT-cat, β-cat, Afadin, and VCL expression in heart tissue lysate from WT (*n*=4) and Y822F (*n*=5) P14 (A) and P28 (B) mice.

Supplemental Figure 1

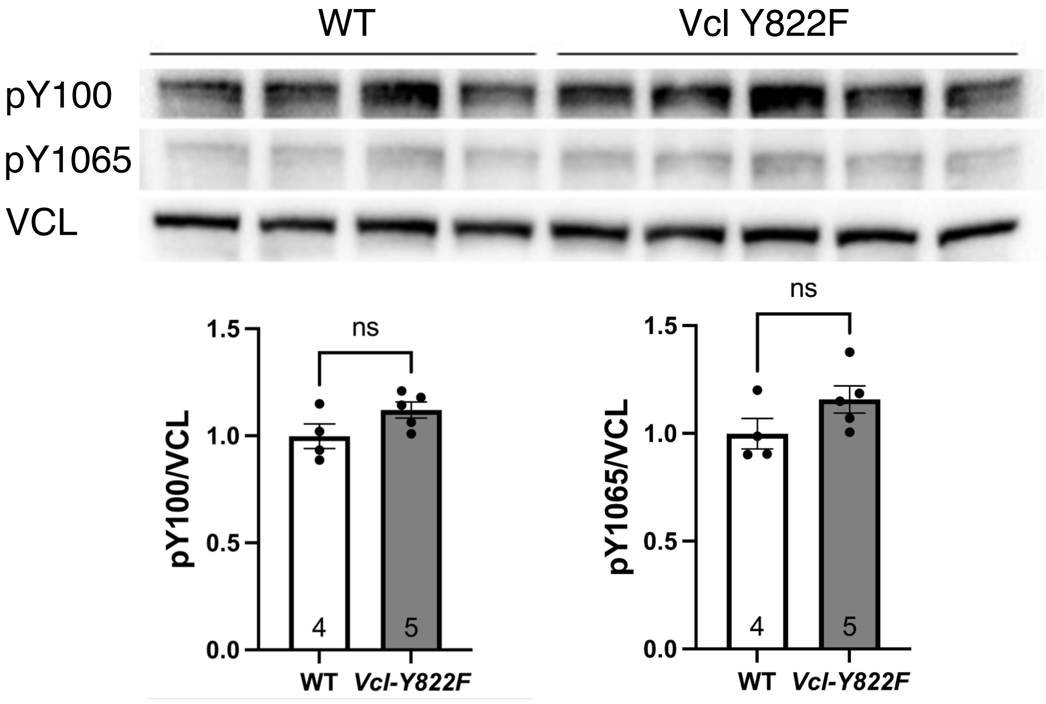


**Supplemental Figure 2. Phosphorylation at VCL Y100 and Y1065 does not change in the *Vcl Y822F* hearts.** Western blots and quantitative analysis of pY100 and pY1065 expression in heart lysates from WT (n=4) and *Vcl Y822F* (n=4) P7 mice.

Supplemental Figure 3

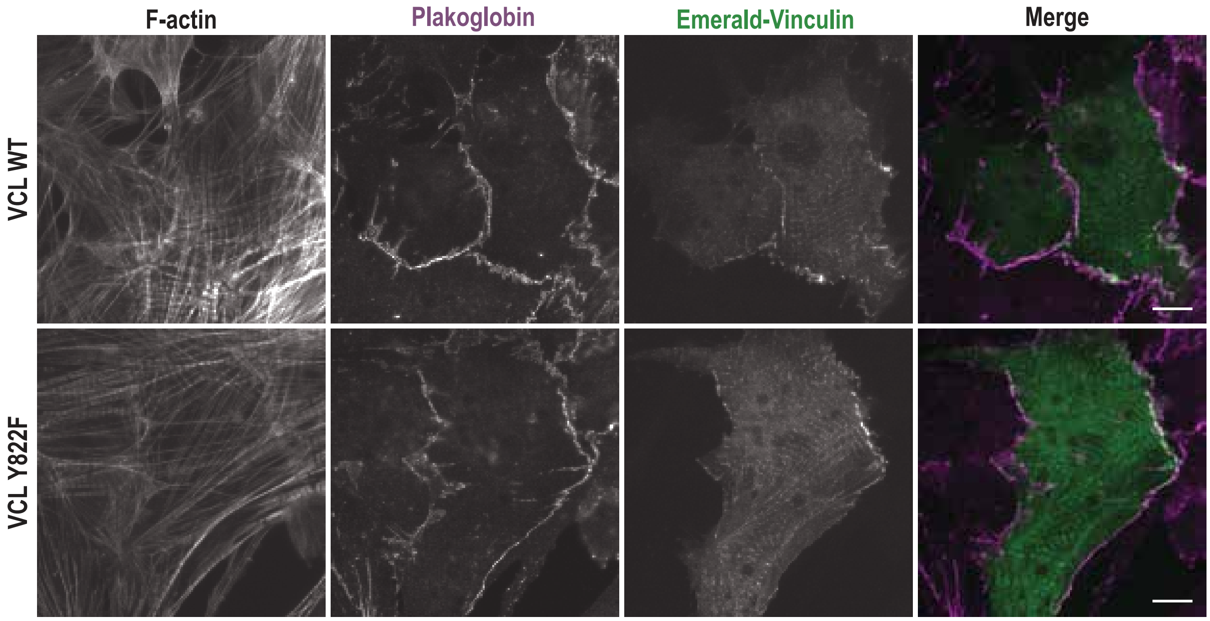


**Supplemental Figure 3. Emerald-VCL WT and Emerald-VCL Y882F localize to FAs and AJs in transfected CMs.** Emerald-tagged VCL WT (top row, green in merge) or VCL Y822F (bottom row, green in merge) were transfected into CMs plated on COL, fixed 48 hrs post-transfection, and stained for F-actin and plakoglobin (magenta in merge). Individual channels and Emerald/plakoglobin merge shown. Scale bar 10 μm.

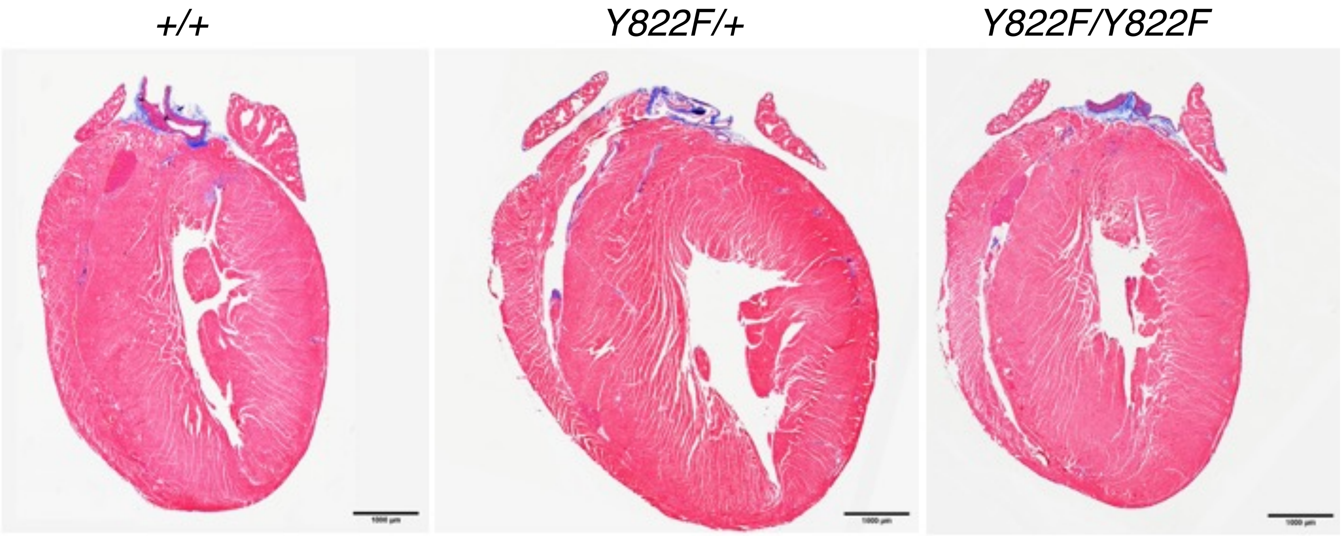


**Supplemental Figure 3. No increase in fibrosis in *Vcl Y822F* hearts.** (A) Representative Masson's Trichrome-stained heart sections from *Vcl +/+*, *Vcl Y822F/+,* and *Vcl Y822F/Y822F* female mice at 40 wks of age.

Supplemental Figure 4.
